# Enhanced triacylglycerol (TAG) and protein accumulation in transgenic diatom *Thalassiosira pseudonana* with altered photosynthetic pigmentation

**DOI:** 10.1101/2020.01.07.897850

**Authors:** Olga Gaidarenko, Daniel P. Yee, Mark Hildebrand

## Abstract

Microalgal productivity in mass cultures is limited by the inefficiency with which available light energy is utilized. In dense cultures, cells closest to the light source absorb more light energy than they can use and dissipate the excess, while light penetrance into the culture is steeply attenuated. Reducing microalgal light harvesting and/or dissipating capacity per cell may improve total light utilization efficiency in mass cultures. In this study, two transgenic lines of the diatom *Thalassiosira pseudonana* with altered photosynthetic pigment content are evaluated with respect to photosynthetic parameters, growth, and macromolecule accumulation. In one line, violaxanthin de-epoxidase-like 2 (VDL2) is overexpressed (OE), resulting in a reduction of the diadinoxanthin cycle pigments, which are involved in light energy dissipation (non-photochemical quenching, NPQ), accompanied by a stoichiometric increase in the light-harvesting pigment fucoxanthin. No differences in the maximum potential quantum yield of photosystem II (Fv/Fm) or light-limited photosynthetic rate (α) were found. However, when adapted to 30 µmol photons m^−2^ sec^−1^, the VDL2 OE maximum relative electron transport rate (rETR_max_) upon exposure to saturating light intensities was 86-95% of wild type (WT). When adapted to 300 µmol photons m^−2^ sec^−1^, VDL2 OE saturated photosynthesis at 62-71% of the light intensity needed to saturate WT (E_k_). NPQ was substantially lower at and below 300 µmol photons m^−2^ sec^−1^. VDL2 OE accumulated up to 3.4 times as much triacylglycerol (TAG) as WT during exponential growth, and up to twice as much protein. Growth in terms of culture density was up to 7% slower. TAG and protein accumulation inversely correlated with NPQ. The second line evaluated was obtained by using antisense RNA to simultaneously silence or knock down (KD) both LUT1-like (LTL) genes, hypothesized to catalyze an intermediate carotenoid biosynthesis step of converting β-carotene to zeaxanthin. Overall reduction of photosynthetic pigment content without altering the relative abundance of individual pigments resulted. No significant differences in photosynthetic parameters compared to WT were found. LTL KD grew at a rate comparable to WT and accumulated up to 40% more TAG during exponential growth, while protein content was reduced by 11-19%. LTL KD cells were elongated and 5-10% smaller than WT, and cultures contained auxospores, indicating stress that may relate to a cell cycle progression defect.

## INTRODUCTION

Microalgae are a promising production platform for sustainable biofuels and other bioproducts. They do not compete with plant crops for arable land and require only light energy and modest nutrient input to synthesize biomolecules of commercial interest and accumulate biomass [Mata et al. 2010]. A major challenge for the economic feasibility of microalgal production is the inefficient utilization of light by microalgal cultures grown at scale [De Mooij et al. 2015]. Microalgae have evolved extensive photosynthetic pigmentation, which confers a competitive advantage in the wild by maximizing a cell’s light absorption capacity in environments where light may be limited, while minimizing the light available to cells below. In dense cultures such as those used for production, this results in a suboptimal distribution of light energy. At high light intensities such as direct sunlight, algae closest to the light source capture more light than the cells are able to utilize. Excess light is wastefully dissipated as heat and fluorescence. Because of the efficiency of light capture, there is a steep attenuation of light penetrance into the culture. Thus, the cells closest to the light source are subject to photosystem-damaging light-induced stress, while the cells deeper in the culture have less light available for photosynthesis [De Mooij et al. 2015]. In theory, reducing cellular light-harvesting and/or dissipation capacity could improve light distribution and therefore productivity in dense cultures, making their cultivation for commercial purposes more cost-efficient. Cells closest to the light source would suffer less light-induced stress, and/or light penetrance would increase. Thus, a greater proportion of the culture would be photosynthetically active [De Mooij et al. 2015].

Numerous efforts have been made in chlorophytes to reduce the size of photoantennae, which serve to capture light energy and funnel it to the photosynthetic reaction centers. This has generally resulted in improved photosynthetic parameters, such as saturation of photosynthesis at higher irradiance, greater light-saturated rates of oxygen evolution on a per-chlorophyll basis [Beckmann et al. 2009, Cazzaniga et al. 2014, Jeong et al. 2017, Kirst et al. 2012a, Kirst et al. 2012b, Mitra and Melis 2008, Nakajima et al. 2001, Shin et al. 2016, Shin et al. 2017], increased quantum yield and lower photoinhibition [Mussgnug et al. 2007]. In some cases, higher maximal culture density [Polle at. al. 2003], faster growth [Mussgnug et al. 2007], and better biomass productivity in laboratory conditions have been reported [Beckman et al. 2009, Shin et al. 2016, Shin et al. 2017]. Several strains that appeared promising based on laboratory performance did not show improved biomass productivity in mass culture conditions simulated in laboratory-scale panel photobioreactors, possibly due to unintended effects of the genetic modifications or higher vulnerability to photodamage [De Mooij et al. 2015]. Cazzaniga et al. [2014], however, reported improved biomass productivity in laboratory conditions as well as in 7 L hanging bag photobioreactors deployed outdoors. As a different strategy, Berteotti et al. [2016] explored downregulation of light energy dissipation through non-photochemical quenching (NPQ) in the chlorophyte *Chlamydomonas reinhardtii*, and found that if it is reduced, but not completely abolished, improved biomass productivity in a small scale photobioreactor results.

Non-chlorophyte eukaryotic microalgae have been largely unexplored with respect to light utilization efficiency improvement through biological modification. Because microalgae are incredibly diverse and have evolved different strategies for interacting with their environment, different taxa may respond to such modifications with varying degrees of success. Diatoms, for example, are brown microalgae belonging to the Stramenopile or Heterokont class, whose light-harvesting and photoprotective strategies differ substantially from chlorophytes [Wilhelm et al. 2006]. Chlorophytes rely predominantly on chlorophylls a and b (Chl a, Chl b) for light capture, the ratio of which adjusts dynamically in response to changes in light intensity. Diatoms use Chl a in association with chlorophyll c (Chl c) and the more abundant carotenoid-derived accessory photosynthetic pigment fucoxanthin (Fx) to capture light energy [Wilhelm et al. 2006]. The ratio of Chl a to Fx, and thus photoantenna size, does not change much with light intensity [Lepetit et al. 2012]. Diatoms appear to rely on their capacity to induce NPQ faster and to a higher extent than chlorophytes when adjusting to short-term irradiance increases and coordinately reduce the abundance of photosynthetic reaction centers and photoantennae during long-term adaptation to higher light [Lepetit et al. 2012]. NPQ in diatoms relies predominantly on the diadinoxanthin (Ddx) cycle that is absent in chlorophytes, wherein the carotenoid derivative Ddx is reversibly converted to diatoxanthin (Dtx) via de-epoxidation when it is necessary to dissipate excess light energy [Lepetit et al. 2012, Wilhelm et al. 2006]. Because diatoms are very promising in terms of biomass productivity and triacylglycerol (TAG, neutral lipid of interest for biofuels) accumulation [Hildebrand et al. 2012], it is intriguing to explore improving their productivity by modulating culture light utilization efficiency. So far, only one such study has been published. A *Cyclotella* sp. strain was obtained through two subsequent rounds of mutagenesis, employing ethylmethylsulfonate and ultraviolet radiation [Huesemann et al. 2009]. Its green color indicated that it had drastically reduced carotenoid abundance, including the main accessory light-harvesting pigment Fx and the photoprotective Ddx cycle pigments. It also had a substantially higher Chl a/Chl c ratio, indicating a smaller photoantenna size. The mutant required higher light intensity to saturate photosynthesis on a per chlorophyll basis but was less stable in culture and had reduced biomass productivity. It was not fully characterized, but the observed lack of fitness may be attributable to too much reduction in carotenoids and thus susceptibility to photodamage and oxidative stress, and possible additional undesirable mutations [Huesemann et al. 2009]. More exploration is necessary to determine the utility of reducing light absorption and/or dissipation in diatoms.

In this study, we evaluate photosynthetic parameters, growth, carbon partitioning, and macromolecule accumulation in two transgenic (TG) lines of the diatom *Thalassiosira pseudonana* in which the abundance of carotenoid biosynthesis enzymes is manipulated. In one line, violaxanthin de-epoxidase-like 2 is overexpressed (VDL2 OE), resulting in a decrease of the photoprotective Ddx cycle pigments (Ddx+Dtx) and a stoichiometric increase in the light-harvesting Fx [Gaidarenko et al. 2020]. In the other line, both copies of LUT1-like (LTL), hypothesized to catalyze an earlier step in the pathway converting β-carotene to zeaxanthin, are simultaneously knocked down (KD), resulting in an overall reduction of total cellular photosynthetic pigment content with conserved ratios of individual pigments [Gaidarenko et al. 2020]. Four clones of each TG line were selected for characterization and compared to wild type (WT) in low light (LL, 30 µmol photons m^−2^ sec^−1^) and high light (HL, 300 µmol photons m^−2^ sec^−1^). There were four experimental culture sets: VDL2 OE vs. WT in LL, VDL2 OE vs. WT in HL, LTL KD vs. WT in LL, and LTL KD vs. WT in HL. Each set was independently acclimated to cultivation conditions and used to obtain samples and data, which were then processed independently of each other. Thus, our findings will be discussed as comparisons within but not between the sets.

## RESULTS

### Photosynthetic Parameters

Photosynthetic parameters were determined for LL and HL-adapted cultures via rapid light-response curves (RLCs) obtained with a pulse amplitude modulation (PAM) fluorometer. This approach involves measuring dynamic rather than steady state responses of a culture to gradual increases in light intensity, and allows, therefore, an assessment of the photosynthetic performance of a culture as it relates to the light intensity it had previously adapted to [Malapascua et al. 2014, Ralph and Gaderman 2005]. The parameters assessed were Fv/Fm (maximum potential quantum yield of photosystem II), α (initial slope of the RLC curve, light-limited photosynthetic rate), rETR_max_ (maximum light-saturated relative electron transport rate), E_k_ (rETR_max_/α, minimum saturating light intensity), and NPQ (non-photochemical quenching, dissipation of photon energy) [Malapascua et al. 2014, Ralph and Gaderman 2005].

No significant differences in Fv/Fm were found between the TG lines and WT in either LL or HL **(Table 1)**. α, rETR_max_, and E_k_ were derived from rETR measurements over a range of irradiance, depicted in **Fig. 1**. LL-adapted VDL2 OE clones had a reduced rETR_max_, calculated to be 86-95% of the WT average (p-value = 0.02) **(Fig. 1A, Table 1)**. However, rETR values did not vary between the TG lines and WT in any tested condition when measured at the irradiance they were adapted to during cultivation **(Fig. 1)**. Additionally, HL-adapted VDL2 OE clones had reduced E_k_values, calculated to saturate photosynthesis at 62-71% of the average WT minimum saturating light intensity (p-value = 0.008) **(Fig. 1B, Table 1)**. No other significant differences in between WT and TG lines were found.

**Fig. 1.**
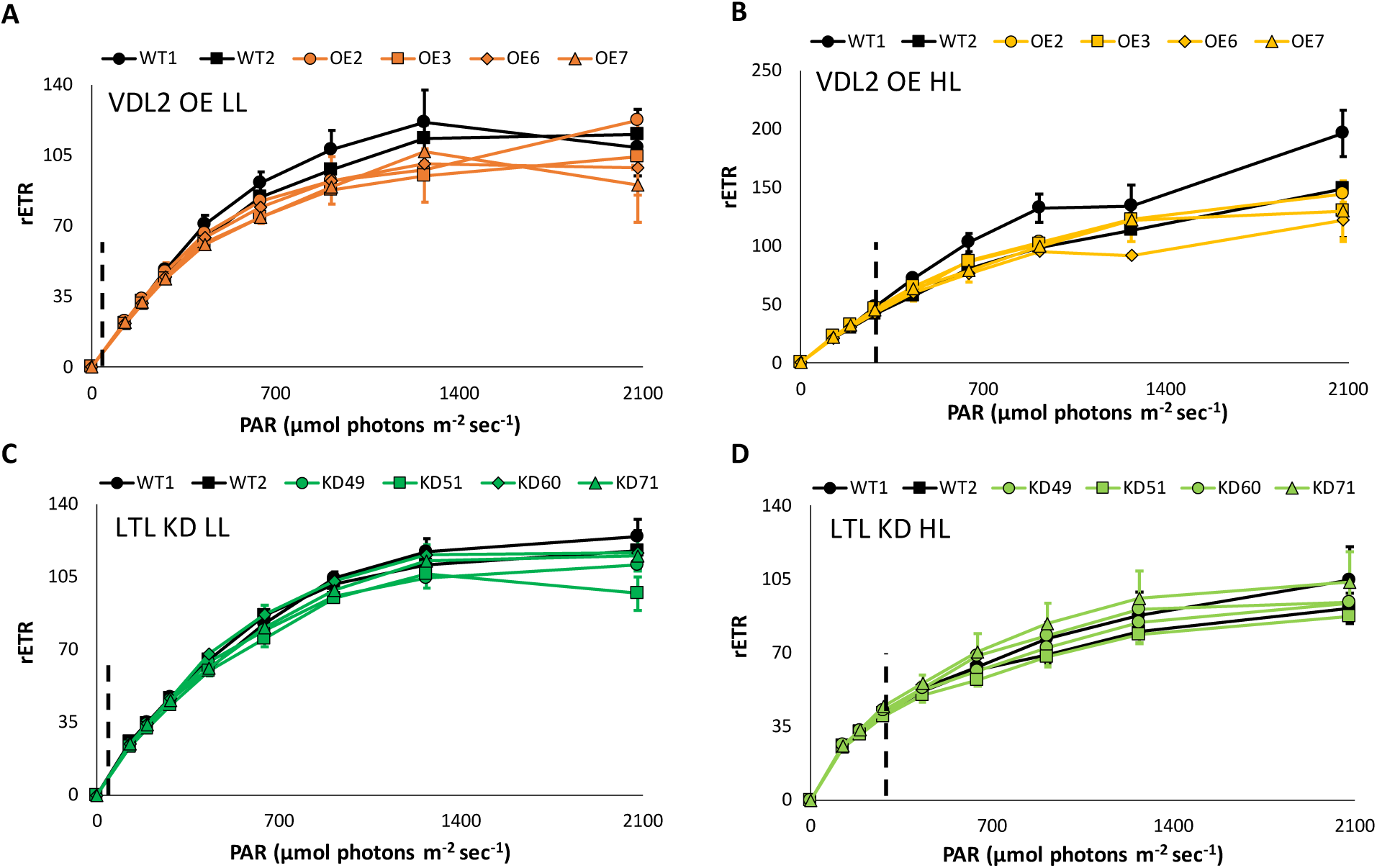
Rapid light curves, relative electron transfer rates (rETR) vs. photosynthetically active radiation (PAR). Vertical dashed lines indicate irradiance used during cultivation. Data are presented as averages of 3-4 replicates ± standard deviation. **A.** 30 µmol photons m^−2^ sec^−1^ (low light - LL)-adapted violaxanthin de-epoxidase-like 2 overexpression (VDL2 OE) clones vs. wild-type (WT); **B.** 300 µmol photons m^−2^ sec^−1^ (high light - HL)-adapted VDL2 OE clones vs. WT; **C.** LL-adapted LUT1-like knockdown (LTL KD) clones vs. WT; **D.** HL-adapted LTL KD clones vs. WT.

**Table 1.** Photosynthetic parameters. Individual violaxanthin de-epoxidase-like 2 overexpression (VDL2 OE) and LUT1-like knockdown (LTL KD) clones are compared to wild-type (WT) cultures in low light (30 µmol photons m^−2^ sec^−1^, LL) and high light (300 µmol photons m^−2^ sec^−1^, HL). Fv/Fm = maximum quantum yield of photosystem II (PSII), α = light-limited photosynthetic rate, rETR_max_ = maximum relative electron transport rate, E_k_ = minimum saturating irradiance.

| | | Fv/Fm | $\alpha$ | $r\text{ETR}_{\text{max}}$ | $E_k$ |
| --- | --- | --- | --- | --- | --- |
| VDL2 OE LL | WT1 | 0.698 ± 0.009 | 0.214 ± 0.032 | 119.0 ± 6.3 | 570.3 ± 99.7 |
| | WT2 | 0.707 ± 0.007 | 0.185 ± 0.021 | 118.3 ± 6.7 | 652.3 ± 112.6 |
| | OE2 | 0.697 ± 0.005 | 0.166 ± 0.006 | 112.3 ± 5.6 | 676.9 ± 34.9 |
| | OE3 | 0.714 ± 0.030 | 0.158 ± 0.010 | 102.1 ± 9.2 | 653.7 ± 96.1 |
| | OE6 | 0.713 ± 0.003 | 0.191 ± 0.031 | 104.3 ± 10.8 | 570.3 ± 162.5 |
| | OE7 | 0.688 ± 0.003 | 0.196 ± 0.002 | 102.9 ± 11.7 | 524.5 ± 64.1 |
|  | *p-value* | *0.9* | *0.3* | ***0.02*** | *0.9* |
| VDL2 OE HL | WT1 | 0.699 ± 0.008 | 0.158 ± 0.008 | 217.1 ± 36.3 | 1368.6 ± 179.4 |
| | WT2 | 0.679 ± 0.007 | 0.135 ± 0.000 | 151.0 ± 5.9 | 1119.9 ± 49.0 |
| | OE2 | 0.685 ± 0.008 | 0.175 ± 0.019 | 146.9 ± 7.8 | 850.5 ± 113.7 |
| | OE3 | 0.680 ± 0.002 | 0.173 ± 0.026 | 133.7 ± 16.5 | 803.6 ± 194.3 |
| | OE6 | 0.678 ± 0.005 | 0.156 ± 0.024 | 115.8 ± 10.4 | 770.8 ± 166.4 |
| | OE7 | 0.685 ± 0.008 | 0.164 ± 0.031 | 136.4 ± 19.4 | 882.8 ± 257.9 |
|  | *p-value* | *0.4* | *0.1* | *0.09* | ***0.008*** |
| LTL KD LL | WT1 | 0.711 ± 0.009 | 0.184 ± 0.018 | 125.4 ± 8.2 | 692.9 ± 95.9 |
| | WT2 | 0.713 ± 0.007 | 0.210 ± 0.007 | 119.9 ± 10.3 | 573.5 ± 67.1 |
| | KD49 | 0.693 ± 0.009 | 0.194 ± 0.022 | 110.7 ± 3.9 | 578.6 ± 68.4 |
| | KD51 | 0.703 ± 0.011 | 0.190 ± 0.010 | 104.5 ± 4.3 | 550.7 ± 33.8 |
| | KD60 | 0.706 ± 0.007 | 0.197 ± 0.020 | 119.9 ± 5.8 | 617.0 ± 95.4 |
| | KD71 | 0.709 ± 0.009 | 0.183 ± 0.017 | 117.3 ± 8.9 | 650.3 ± 111.5 |
|  | *p-value* | *0.2* | *0.6* | *0.2* | *0.5* |
| LTL KD HL | WT1 | 0.639 ± 0.006 | 0.141 ± 0.007 | 97.9 ± 17.9 | 702.8 ± 150.5 |
| | WT2 | 0.625 ± 0.002 | 0.147 ± 0.003 | 85.0 ± 5.0 | 581.4 ± 44.2 |
| | KD49 | 0.609 ± 0.004 | 0.158 ± 0.008 | 81.8 ± 11.2 | 517.4 ± 55.4 |
| | KD51 | 0.628 ± 0.009 | 0.153 ± 0.016 | 72.2 ± 11.0 | 483.1 ± 113.4 |
| | KD60 | 0.625 ± 0.004 | 0.172 ± 0.023 | 92.3 ± 10.5 | 554.6 ± 131.9 |
| | KD71 | 0.647 ± 0.027 | 0.154 ± 0.014 | 92.9 ± 15.1 | 617.0 ± 138.9 |
|  | *p-value* | *0.7* | *0.08* | *0.5* | *0.2* |

RLC-derived NPQ values are depicted in **Fig. 2**. A high degree of variability between different TG clones and WT cultures was observed, and no claims about statistically significant differences consistent between all TG clones within any condition could thus be made, with one exception. At lower irradiance values, the HL-adapted VDL2 OE clones had substantially less NPQ than WT **(Fig. 2B)**. At 125 µmol photons m^−2^ sec^−1^, VDL2 OE clone 6 had 53% of the average WT NPQ, while the other three clones were at 1-8% of the WT average (p-value = 0.01). At 191 µmol photons m^−2^ sec^−1^, VDL2 OE clone 6 measured at 64% of the WT average NPQ, while the other three clones had 16-18% of the WT average (p-value = 0.02). At 282 µmol photons m^−2^ sec^−1^, VDL2 OE clone 6 had 82% of the average WT NPQ, while the other three clones were at 27-36% (p-value = 0.08 with clone 6, and 0.03 without).

**Fig. 2.**
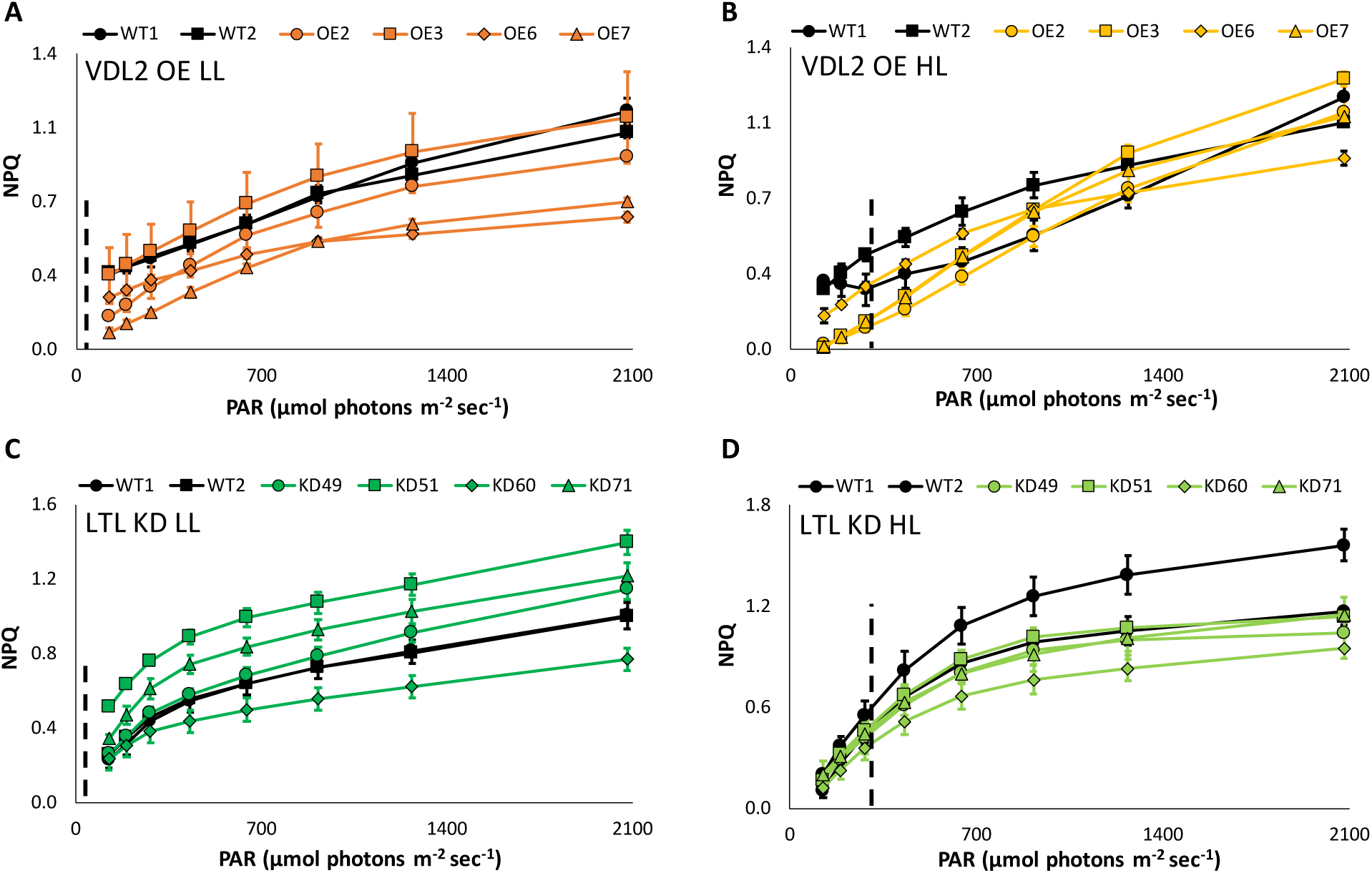
Rapid light curve-derived non-photosynthetic quenching (NPQ) values vs. photosynthetically active radiation (PAR). Vertical dashed lines indicate irradiance used during cultivation. Data are presented as averages of 3-4 replicates ± standard deviation. **A.** 30 µmol photons m^−2^ sec^−1^ (low light - LL)-adapted violaxanthin de-epoxidase-like 2 overexpression (VDL2 OE) clones vs. wild-type (WT); **B.** 300 µmol photons m^−2^ sec^−1^ (high light - HL)-adapted VDL2 OE clones vs. WT; **C.** LL-adapted LUT1-like knockdown (LTL KD) clones vs. WT; **D.** HL-adapted LTL KD clones vs. WT.

### Morphology, Cell and Chloroplast Size

No morphological differences were observed between WT and VDL2 OE clones. LTL KD clones, on the other hand, were elongated and formed numerous auxospores, indicating sexual reproduction **(Fig. 3A, B)**. The elongation phenotype and auxospore formation were more prominent in HL, but observable in LL as well. Incubating with PDMPO, a fluorescent dye that incorporates into newly synthesized silica, confirmed that the elongated cells were single cells, not multiple cells that have failed to separate, as no cell wall could be observed within the cells **(Fig. 3C, D)**. The LTL KD cells with a high degree of elongation appeared to have numerous chloroplasts distributed throughout **(Fig. 3E, F)**.

**Fig. 3.**
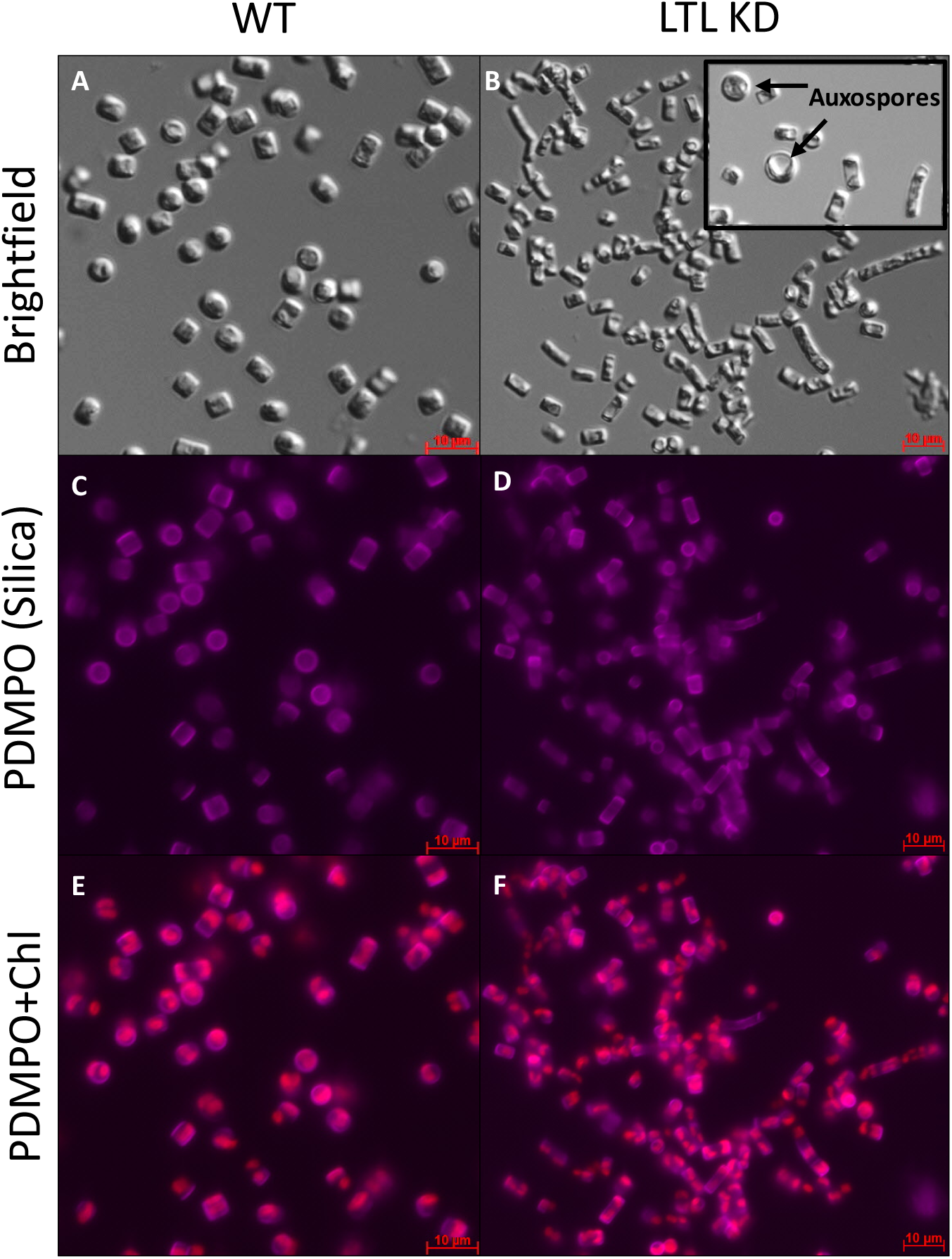
Wild-type (WT) (A, C, E) and LUT1-like knockdown (LTL KD) (B, D, F) cultures, cultivated at 300 µmol photons m^−2^ sec^−1^. **A, B.** Brightfield. Arrows indicate auxospores; **C, D.** PDMPO staining for silica; **E, F.** PDMPO and chlorophyll fluorescence (Chl).

The average cell area was not significantly different between WT and VDL2 OE clones but was reduced in LTL KD clones in HL, measuring at 90-95% of the WT average (p-value = 0.03) **(Table 2)**. The chloroplast to cell area ratio in LTL KD clones did not significantly differ from WT in LL or HL **(Table 2)**. In HL-adapted VDL2 OE clones, the chloroplast to cell area ratio was reduced to 93-96% of the WT average (p-value = 0.02).

**Table 2.**
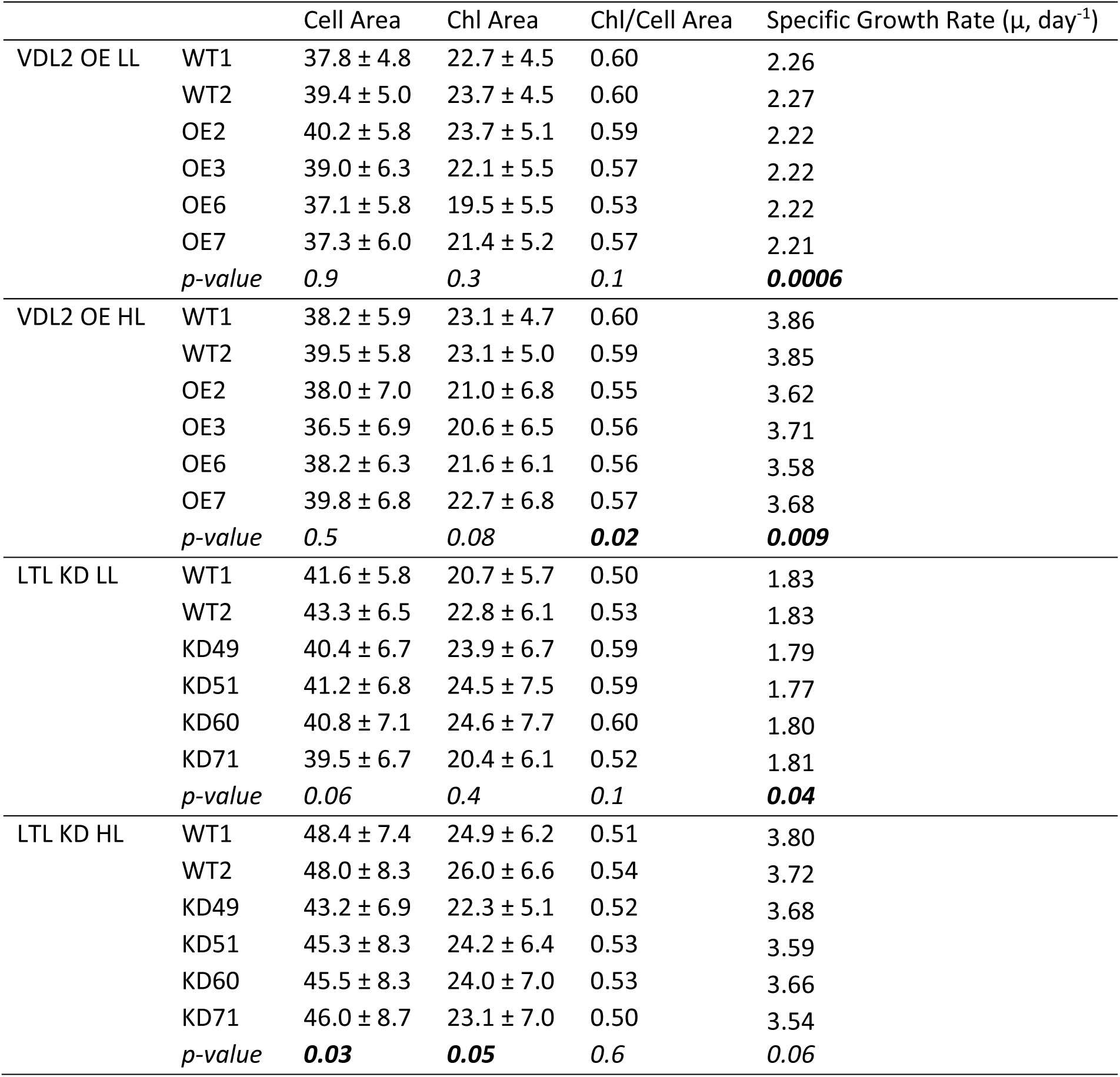
Average cell and chloroplast (Chl) area measurements, their ratios, and specific growth rates. Individual violaxanthin de-epoxidase-like 2 overexpression (VDL2 OE) and LUT1-like knockdown (LTL KD) clones are compared to wild-type (WT) cultures in low light (30 µmol photons m^−2^ sec^−1^, LL) and high light (300 µmol photons m^−2^ sec^−1^, HL).

| | | Cell Area | Chl Area | Chl/Cell Area | Specific Growth Rate ($\mu$, day<sup>-1</sup>) |
| --- | --- | --- | --- | --- | --- |
| VDL2 OE LL | WT1 | 37.8 ± 4.8 | 22.7 ± 4.5 | 0.60 | 2.26 |
| | WT2 | 39.4 ± 5.0 | 23.7 ± 4.5 | 0.60 | 2.27 |
| | OE2 | 40.2 ± 5.8 | 23.7 ± 5.1 | 0.59 | 2.22 |
| | OE3 | 39.0 ± 6.3 | 22.1 ± 5.5 | 0.57 | 2.22 |
| | OE6 | 37.1 ± 5.8 | 19.5 ± 5.5 | 0.53 | 2.22 |
| | OE7 | 37.3 ± 6.0 | 21.4 ± 5.2 | 0.57 | 2.21 |
|  | *p-value* | 0.9 | 0.3 | 0.1 | **0.0006** |
| VDL2 OE HL | WT1 | 38.2 ± 5.9 | 23.1 ± 4.7 | 0.60 | 3.86 |
| | WT2 | 39.5 ± 5.8 | 23.1 ± 5.0 | 0.59 | 3.85 |
| | OE2 | 38.0 ± 7.0 | 21.0 ± 6.8 | 0.55 | 3.62 |
| | OE3 | 36.5 ± 6.9 | 20.6 ± 6.5 | 0.56 | 3.71 |
| | OE6 | 38.2 ± 6.3 | 21.6 ± 6.1 | 0.56 | 3.58 |
| | OE7 | 39.8 ± 6.8 | 22.7 ± 6.8 | 0.57 | 3.68 |
|  | *p-value* | 0.5 | 0.08 | **0.02** | **0.009** |
| LTL KD LL | WT1 | 41.6 ± 5.8 | 20.7 ± 5.7 | 0.50 | 1.83 |
| | WT2 | 43.3 ± 6.5 | 22.8 ± 6.1 | 0.53 | 1.83 |
| | KD49 | 40.4 ± 6.7 | 23.9 ± 6.7 | 0.59 | 1.79 |
| | KD51 | 41.2 ± 6.8 | 24.5 ± 7.5 | 0.59 | 1.77 |
| | KD60 | 40.8 ± 7.1 | 24.6 ± 7.7 | 0.60 | 1.80 |
| | KD71 | 39.5 ± 6.7 | 20.4 ± 6.1 | 0.52 | 1.81 |
|  | *p-value* | 0.06 | 0.4 | 0.1 | **0.04** |
| LTL KD HL | WT1 | 48.4 ± 7.4 | 24.9 ± 6.2 | 0.51 | 3.80 |
| | WT2 | 48.0 ± 8.3 | 26.0 ± 6.6 | 0.54 | 3.72 |
| | KD49 | 43.2 ± 6.9 | 22.3 ± 5.1 | 0.52 | 3.68 |
| | KD51 | 45.3 ± 8.3 | 24.2 ± 6.4 | 0.53 | 3.59 |
| | KD60 | 45.5 ± 8.3 | 24.0 ± 7.0 | 0.53 | 3.66 |
| | KD71 | 46.0 ± 8.7 | 23.1 ± 7.0 | 0.50 | 3.54 |
|  | *p-value* | **0.03** | **0.05** | 0.6 | 0.06 |

### Growth Rates and Maximal Culture Density

During exponential growth in LL, the specific growth rate of the VDL2 OE clones was approximately 98% of the WT average (p-value = 0.0006) **(Table 2)**. When light-limiting culture densities were reached around day 7, the disparity between the KD clones and WT cultures became more pronounced **(Fig. 4A)**. The stationary phase culture densities reached by the VDL2 OE clones in LL were 72-88% of the WT average (p = 0.02).

**Fig. 4.**
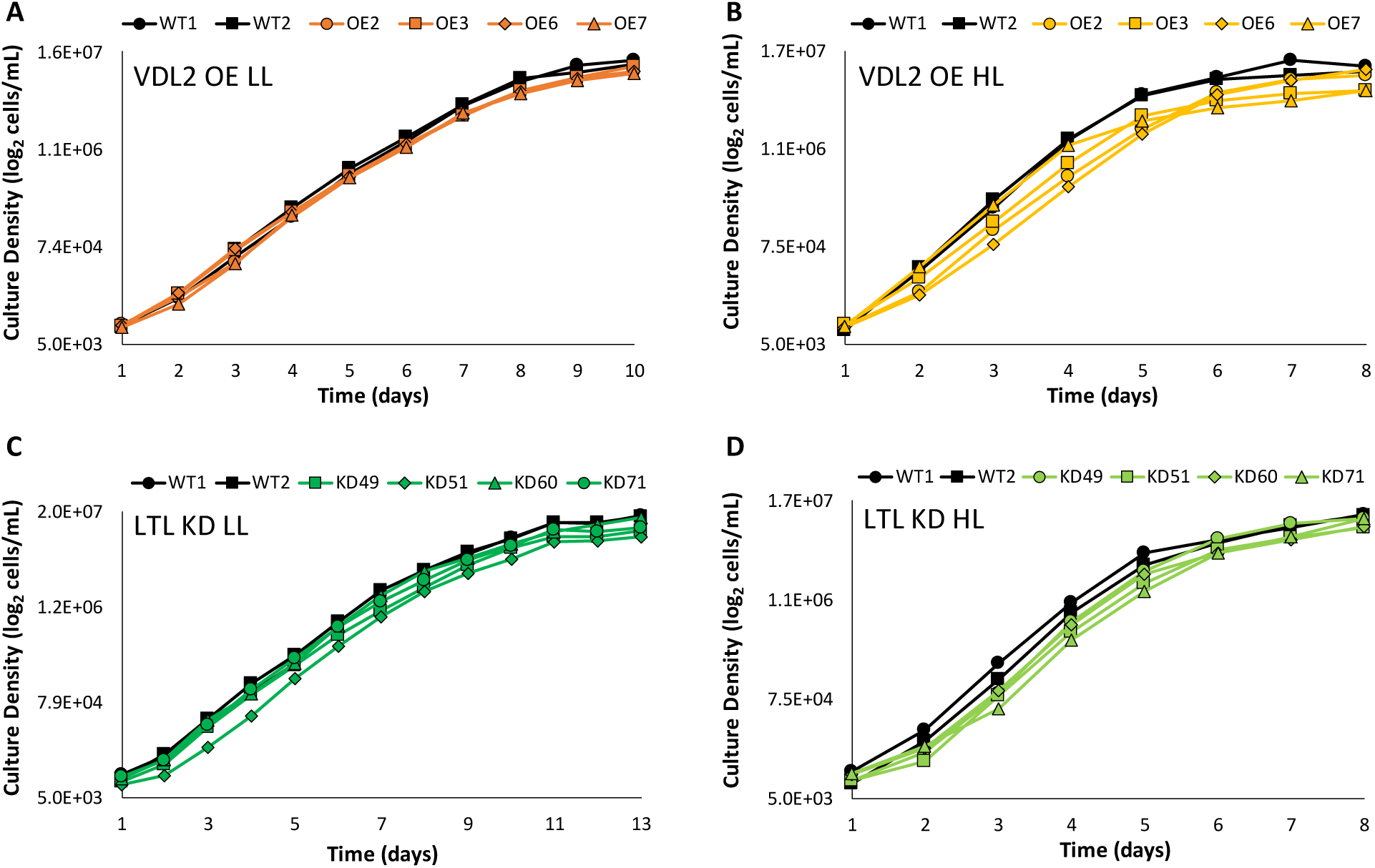
Growth curves. **A.** 30 µmol photons m^−2^ sec^−1^ (low light - LL)-adapted violaxanthin de-epoxidase-like 2 overexpression (VDL2 OE) clones vs. wild-type (WT); **B.** 300 µmol photons m^−2^ sec^−1^ (high light - HL)-adapted VDL2 OE clones vs. WT; **C.** LL-adapted LUT1-like knockdown (LTL KD) clones vs. WT; **D.** HL-adapted LTL KD clones vs. WT.

In HL, VDL2 OE clones also experienced slowing **(Fig. 4B)**. During exponential growth, the specific growth rate of the VDL2 OE clones was 93-96% of the WT average (p-value = 0.009) **(Table 2)**. At stationary phase, HL clones VDL2 OE3 and OE5 reached 54% of the average WT culture density, OE2 was at 84% of WT, and OE6 did not differ from WT. Interestingly, all the HL-adapted VDL2 OE clones were substantially lower in culture density than WT during day 5 (34-57% of the WT average, p-value = 0.002) and day 6 (45-69% of the WT average, p-value = 0.007), as the light-limited cultures were transitioning to stationary phase **(Fig. 4B)**.

The specific growth rates of the LTL KD clones in LL were 97-99% of the WT average (p-value = 0.04), and not significantly different in HL **(Table 2)**. The increased disparity from WT during the exponential to stationary phase transition that was seen in the VDL2 OE lines was not observed in the LTL KD lines **(Fig. 4)**. Stationary phase culture densities of the LTL KD clones did not significantly differ from WT.

### Lipid, Protein, Carbohydrate, and Photosynthetic Pigment Content

The cellular abundance of TAG (quantified by BODIPY fluorescence), proteins, carbohydrates, and photosynthetic pigments was assessed during exponential growth.

The TAG content of VDL2 OE clones was found to be 21-81% greater than the WT average (p-value = 0.05) in LL, and 2-3.4 times the WT average in HL (p-value = 0.04) (Fig. 5A, B). The LTL KD TAG content did not significantly differ from WT in LL. In HL, LTL KD clone 49 had TAG content similar to WT, while the other 3 clones had 25-40% more (Fig. 5C, D). When normalized to average cell area, LTL KD TAG content, including clone 49, was 17-46% greater than the WT average in HL (p-value = 0.03) **(Fig. S1A)**.

**Fig. 5.**
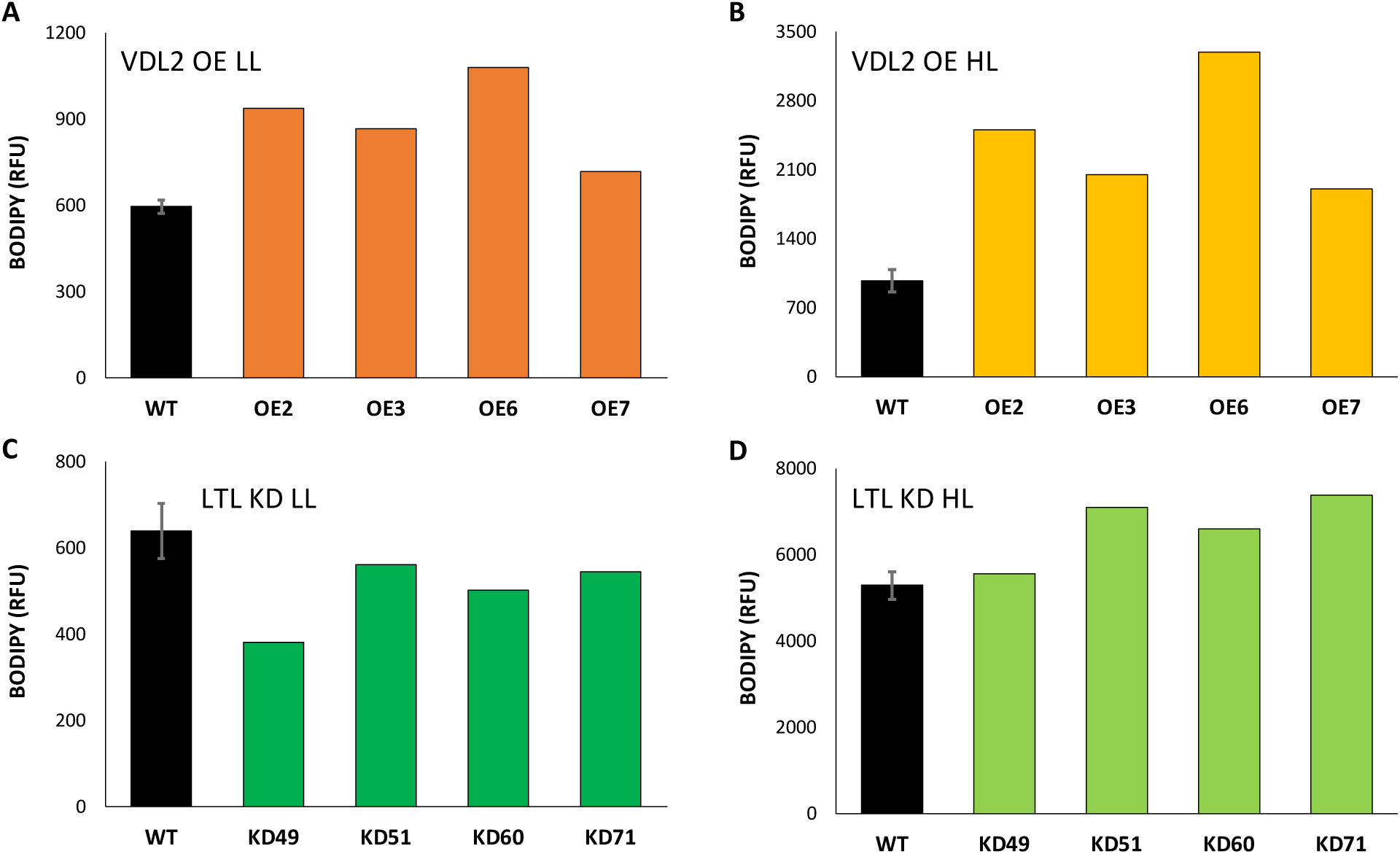
Average BODIPY fluorescence (relative fluorescence units – RFU). Wild-type (WT) data are presented as averages of two independent cultures ± standard deviation. **A.** 30 µmol photons m^−2^ sec^−1^ (low light - LL)-adapted violaxanthin de-epoxidase-like 2 overexpression (VDL2 OE) clones vs. wild-type (WT); **B.** 300 µmol photons m^−2^ sec^−1^ (high light - HL)-adapted VDL2 OE clones vs. WT; **C.** LL-adapted LUT1-like knockdown (LTL KD) clones vs. WT; **D.** HL-adapted LTL KD clones vs. WT.

No significant differences were found when comparing total cellular protein between TG lines and WT in LL **(Fig. 6A, C)**. HL-adapted VDL2 OE clones contained approximately 1.5-2 times as much protein as WT (p-value = 0.02) **(Fig. 6B)**. HL-adapted LTL KD clones had 81-89% of the average WT total cellular protein content (p-value = 0.01) **(Fig. 6D)**. When normalized to average cell area, total cellular protein content did not significantly differ between HL-adapted LTL KD clones and WT **(Fig. S1B)**.

**Fig. 6.**
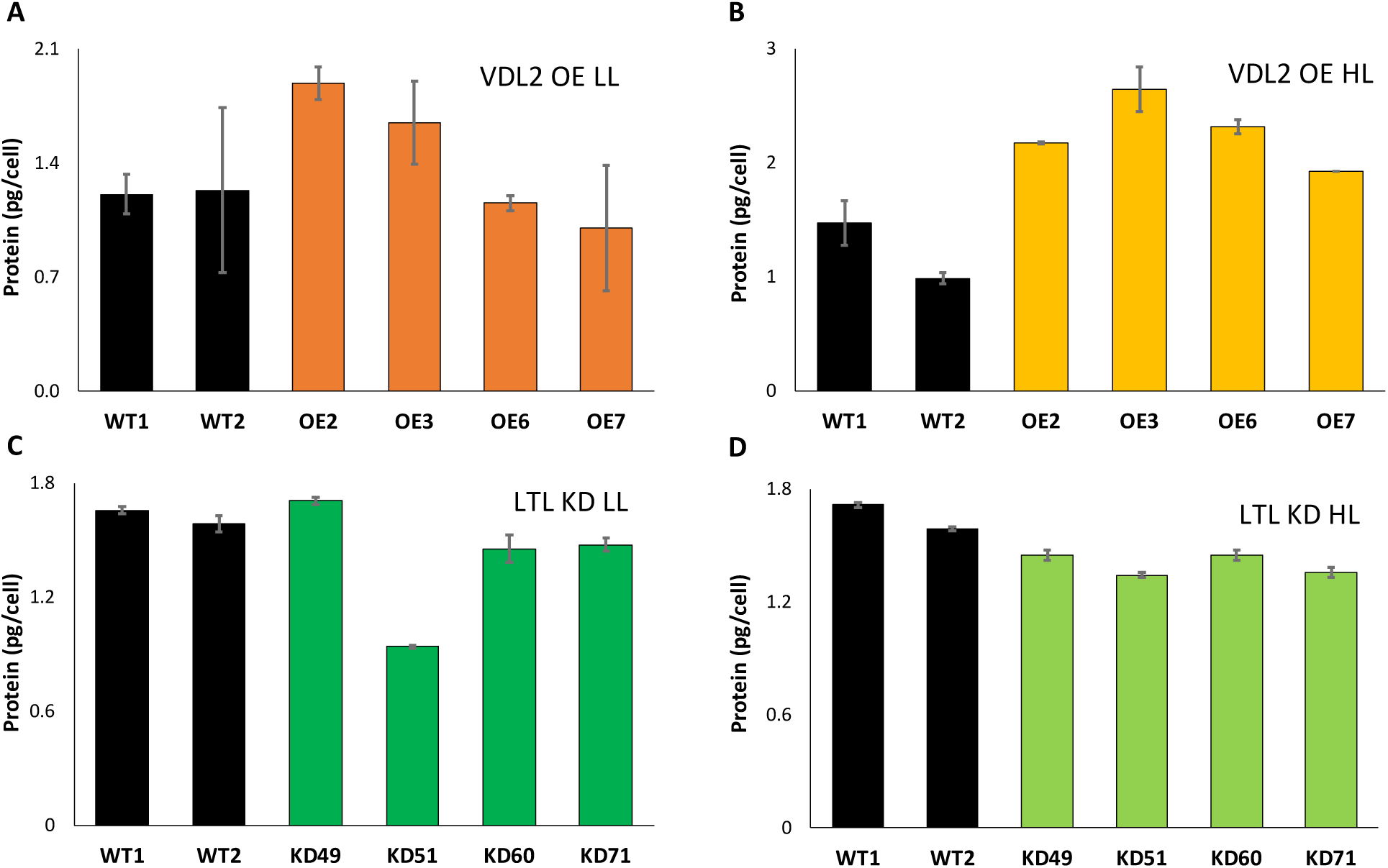
Average protein abundance per cell. Data are presented as averages of two technical replicates ± standard deviation. **A.** 30 µmol photons m^−2^ sec^−1^ (low light - LL)-adapted violaxanthin de-epoxidase-like 2 overexpression (VDL2 OE) clones vs. wild-type (WT); **B.** 300 µmol photons m^−2^ sec^−1^ (high light - HL)-adapted VDL2 OE clones vs. WT; **C.** LL-adapted LUT1-like knockdown (LTL KD) clones vs. WT; **D.** HL-adapted LTL KD clones vs. WT.

No significant differences in total carbohydrate content were found between TG lines and WT, in either LL or HL **(Fig. S2)**. Cell area-normalized carbohydrate content also did not differ significantly between LTL KD clones and WT in HL **(Fig. S1C)**.

Total cellular photosynthetic pigment (Tot) content did not significantly differ between VDL2 OE clones and WT [Gaidarenko et al. 2020]. HL-adapted LTL KD clones had 60-76% of the average WT Tot in HL (p-value = 0.009) [Gaidarenko et al. 2020]. In HL, cell area-normalized Tot was reduced in LTL KD clones to 63-81% of the WT average (p-value = 0.02). In LL-adapted LTL KD clones, a trend in Tot reduction with respect to WT was observable, but not statistically significant [Gaidarenko et al. 2020].

### Inverse Relationship Between NPQ and TAG, Protein Content

NPQ measured at 282 µmol photons m^−2^ sec^−1^, closest to the cultivation conditions for the HL-adapted cultures, was plotted against BODIPY fluorescence and protein content per cell for HL-adapted VDL2 OE clones vs. WT **(Fig. 7)**. The Pearson correlation coefficient (R), where 0 signifies no correlation and −1 means that there is a perfect negative correlation, was −0.4 for BODIPY and −0.7 for protein.

**Fig. 7.**
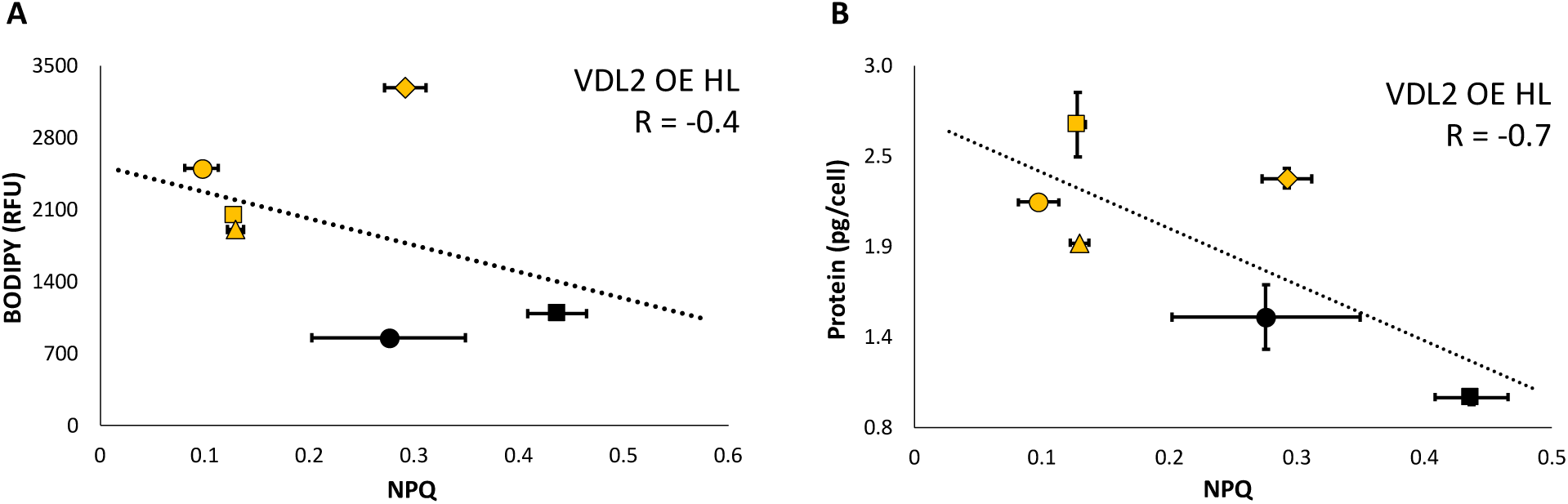
Relationship between non-photochemical quenching (NPQ) measured at 282 µmol photons m^−2^ sec^−1^ and **A.** BODIPY fluorescence (relative fluorescence units, RFU), or **B.** Protein content per cell, for cultures cultivated at 300 µmol photons m^−2^ sec^−1^. Pearson correlation coefficient (R) is indicated on the plots. NPQ values are presented as averages of 3-4 replicates ± standard deviation. Wild-type (WT) cultures are represented by black circles (WT1) and squares (WT2). Violaxanthin de-epoxidase-like 2 overexpression (VDL2 OE) clones are represented by orange circles (VDL2 OE2), squares (VDL2 OE3), rhombuses (VDL2 OE6), and triangles (VDL 2 OE7).

## DISCUSSION

### Photosynthesis

In VDL2 OE clones, Tot did not exhibit a trend with respect to WT, and the ratios of Chl a, Chl c, and β-carotene to Tot were unchanged [Gaidarenko et al. 2020]. This indicates that the stoichiometric increase in Fx/Tot at the expense of (Ddx+Dtx)/Tot did not affect the ratios of photosynthetic reaction centers to photoantennae. Rather, Fx likely replaced some of what would have been Ddx cycle pigments, in one or more of their distinct pools. Some are bound to photoantenna proteins, while others are dissolved in the lipid shield around photoantennae [Lepetit et al. 2010].

Photosynthesis in VDL2 OE clones was not negatively impacted in LL or HL, as indicated by no change in Fv/Fm, α, and rETR measured at the irradiance to which the cultures were adapted with respect to WT **(Table 1, Fig. 1A, B)**. The lowered rETR_max_ in LL-adapted VDL2 OE clones means that in the short term, LL-adapted VDL2 OE clones would have an impairment in photosynthetic electron transfer through photosystem II compared to WT upon being exposed to saturating irradiance **(Table 1, Fig. 1A)**. This is not the case with HL-adapted VDL2 OE clones, the rETR_max_ of which did not significantly differ from WT **(Table 1, Fig. 1B)**. Photosynthetic electron transport of the HL-adapted VDL2 OE clones, however, would, in the short term, saturate at a lower irradiance than in WT, as indicated by lower E_k_ values **(Table 1, Fig. 1B)**.

As discussed previously [Gaidarenko et al. 2020], the overall photosynthetic pigment reduction in LTL KD clones may be attributable to destabilization of light-harvesting complexes caused by Fx insufficiency, as carotenoids are known to be crucial for their assembly and stabilization [Moskalenko and Karapetyan 1996, Santabarbara et al. 2013]. Nevertheless, in comparison to WT, photosynthetic parameters as indicated by Fv/Fm, α, and rETR at the irradiance to which the cultures were adapted were not adversely affected in the LTL KD clones in LL or HL **(Table 1, Fig. 1C, D)**. In contrast to the VDL2 OE clones, short-term adaptation to increases in irradiance was also not impaired, as demonstrated by rETR_max_ and E_k_ values not differing between the LTL KD clones and WT **(Table 1, Fig. 1)**.

The high degree of variance between different TG clones in NPQ responses over a range of irradiance, especially in the LL-adapted state in which WT cultures had a highly replicable response **(Fig. 2)**, could be attributable to potential inter-clonal differences in adaptation to the altered photosynthetic pigment content. The substantially reduced NPQ in HL-adapted VDL2 OE lines at lower irradiance levels **(Fig. 2B)** may be attributed to the reduced Ddx cycle pigment abundance, as the largest part of NPQ in diatoms requires the presence of Dtx [Lepetit et al. 2012]. A strong correlation between Ddx cycle pigment abundance and NPQ was observed for VDL2 OE clones and WT cultures in LL (R = 0.7) and HL (R = 0.8) **(Fig. S3)**. The extent of NPQ induction depends on the light adaptation state and incident irradiance, and it appears that despite the reduced abundance of Ddx cycle pigments, HL-adapted VDL2 OE clones were able to induce WT-equivalent NPQ levels at higher irradiance levels **(Fig. 2B)** [Lepetit et al. 2012].

### Auxospore Formation in LTL KD

Cell elongation and frequent auxospore formation, as observed for the LTL KD clones **(Fig. 3)**, are both markers of stress. Under typical laboratory conditions, auxospore formation, which indicates sexual reproduction, is rarely if ever observed in *T. pseudonana* [Moore et al. 2017]. In some diatom species, sexual reproduction serves as a way to reconstitute cell size, as it diminishes with every division. However, *T. pseudonana* appears to maintain a relatively constant cell size. Sexual reproduction in diatoms may also be triggered by growth stress that leads to cell cycle arrest, such as nutrient depletion and oxidative stress [Moore et al. 2017]. Because the LTL KD clones were grown in the same nutrient-replete media as WT, which did not form abundant auxospores, nutrient depletion is not a likely cause of the observed sexual reproduction in LTL KD clones. Oxidative stress is also unlikely, since it would be expected to lead to reduced Fv/Fm [Jallet et al. 2016], which was not observed **(Table 1)**. A small portion of LTL KD cultures consisted of cells that were smaller than typical, and it is possible that sexual reproduction was induced in those cells to restore size. However, stress evidenced by elongation **(Fig. 3)** was present in the majority of LTL KD cells, and we hypothesize that it was a contributor to auxospore formation. The elongated phenotype has been documented in *T. pseudonana* subject to various stresses that impede cell cycle progression, such as copper toxicity and limitation in silica or selenium. It has not been observed in response to nitrate or phosphorus limitation and is thus not a universal stress response [Davis et al. 2005].

The cause of the stress in our study is not clear. We hypothesize that it may relate to a defect in chloroplast division, stemming from light-harvesting assembly impairment and destabilization by Fx deficiency in LTL KD clones. Microalgae coordinate cell and chloroplast division to ensure that both daughter cells have chloroplasts upon cytokinesis [Sumiya et al. 2016]. Depending on the timing, an arrest in chloroplast division may cause cell cycle arrest, or chloroplasts may continue dividing without concomitant cell cycle progression if cells are arrested in S-phase, resulting in numerous chloroplasts per cell [Sumiya et al. 2016]. A deregulated coordination between chloroplast division and cell cycle progression in the LTL KD clones could thus account for cell elongation and auxospore formation, which may result from a cell cycle progression defect, as well as for the overaccumulation of chloroplasts observed in some of the highly elongated cells **(Fig. 3)**.

### Growth, Carbon Partitioning, and Macromolecule Accumulation

Exponential growth rates for VDL2 OE clones were up to 7% lower than in WT in HL and 2% lower in LL, and average cell area was not statistically different **(Table 2)**. However, VDL2 OE clones accumulated substantially more TAG per cell than WT in LL and HL **(Fig. 5A, B)**, and HL-adapted VDL2 OE clones also accumulated more protein per cell than WT **(Fig. 6B)**. This suggests that VDL2 OE clones may have been fixing more carbon than WT, and preferentially storing it as TAG (and producing more protein in HL), rather than using it to fuel faster growth or storing it as carbohydrate **(Fig. S2B)**. Thus, excess fixed carbon was diverted to glycolysis, which eventually feeds into TAG and amino acid biosynthesis, rather than gluconeogenesis, which is more costly in terms of energy results in carbohydrate biosynthesis [Smith et al. 2012].

Because NPQ values were derived from an RLC curve on dark-adapted cultures, the measurements at the light intensity closest to the irradiance used during cultivation are estimates of the NPQ response in cultures under cultivation conditions. We observed an inverse correlation between NPQ at the cultivation irradiance and TAG as well as protein levels for HL-adapted VDL2 OE clones and WT cultures **(Fig. 7)**. We suggest that diminished NPQ in cultivation conditions allowed the VDL2 OE clones to utilize more photons for carbon fixation, which was then used to synthesize additional TAG and protein.

HL-adapted LTL KD clones had higher TAG than WT **(Figs. 5D, S1A)**, but no significant difference in NPQ **(Fig. 2D)**. We suggest that in this case, enhanced TAG accumulation may have occurred due to the observed cellular stress, with some of the fixed carbon stored as TAG instead of being used to fuel growth. Additionally, since the photosynthetic pigment abundance was reduced, carbon that would have been used for their synthesis may have been repurposed for TAG. Improved light penetrance into the culture due to reduced cellular pigmentation in LTL KD clones may have also contributed to increased carbon fixation and storage as TAG.

### Implications for Diatom Strain Improvement

Our results indicate that reducing the photoprotective Ddx cycle pigments without reducing light-harvesting pigmentation may be a promising strategy for improving TAG and protein productivity in diatoms. Ddx cycle pigments are necessary for NPQ induction and preventing photoinhibition due to excess absorbed light energy and resultant oxidative stress [Lepetit et al. 2010]. However, diatoms may accumulate them in excess of what is needed to protect cells without unnecessarily reducing the amount of light energy available for photosynthesis. A similar concept has been reported by Berteotti et al. [2016], who found that downregulating but not completely abolishing NPQ in the chlorophyte *Chlamydomonas reinhardtii* improved biomass productivity in laboratory conditions.

The ability to enhance TAG production without adversely affecting growth is an important goal for advancing biofuels, as currently their commercialization is stymied by production inefficiency [Trentacoste et al. 2013]. Our VDL2 OE clones suffered somewhat from slower growth with respect to culture density compared to WT. Strain performance in laboratory conditions does not always relate to productivity in a production setting [Gaidarenko et al. 2019]. Thus, it will be important to evaluate VDL2 OE performance in production-relevant conditions. Biomass productivity should also be assessed, as VDL2 OE may have an advantage over WT due to fixing more carbon. If so, slower growth in terms of culture density would be less of a concern. Additionally, the enhanced protein content in VDL2 OE could be used as a high-value co-product for applications such as animal feed, helping offset the costs of TAG production [Moreno-Garcia et al. 2017].

There may be ways to improve upon the VDL2 OE phenotype. The excess Fx in VDL2 OE clones may cause shading and reduce the light utilization efficiency in cultures. One strategy may be to reduce Ddx cycle pigments without changing the amount of Fx. As previously described [Gaidarenko et al. 2019], Ddx cycle pigments appear to be regulated by light intensity, while Fx levels are also affected by cell cycle progression. Since they are both end products of the same biosynthetic pathway, this suggests differential regulation [Gaidarenko et al. 2020] that could be exploited for strain improvement. Another useful approach may be to target proteins involved in NPQ, such as LHCX3 [Hao et al. 2018].

Reducing overall photosynthetic pigment content may also be considered for TAG productivity improvement in diatoms. Although the LTL KD clones had a slightly reduced protein content and did not accumulate as much additional TAG as VDL2 OE clones did, there was an approximately 25-40% improvement over WT in three out of the four clones **(Fig. 5D)**. The disadvantage of the LTL KD clones is the apparent stress experienced by the cells. Nevertheless, it would be interesting to evaluate the performance of LTL KD clones in production conditions to assess the utility of overall photosynthetic pigment reduction for improving productivity in diatoms.

## METHODS

### Cultivation, Sampling, and Growth Curves

WT and TG *T. pseudonana* cultures were cultivated at either 30 or 300 µmol photons m^−2^ sec^−1^ (natural white LED lighting, superbrightleds.com, NFLS-NW300X3-WHT-LC2), using a 12:12 light:dark regime, at 18°C. 50 mL cultures in Erlenmeyer flasks were maintained in Artificial Sea Water (ASW) medium [Darley and Volcani 1969] with rapid stirring. Each experimental set included 2 WT cultures and 4 TG clones. After inoculation, the cultures were grown to 1-3×10^6^ cells/mL, then allowed to adapt to the cultivation conditions by daily dilutions that maintained exponential growth with culture density under 2.5×10^6^ cells/mL for a minimum of 2 weeks prior to sampling. Cultures were rotated between stir plates each day to minimize any position-specific differences and transferred to clean flasks once a week. Sampling for protein content, carbohydrate content, cell/chloroplast dimensions and lipid content, and photosynthetic measurements was performed on separate days, within the first two hours of the light period. After sampling was completed, the cultures were inoculated into fresh 50 mL of ASW for growth curves at approximately 8-11×10^3^ cells/mL. Cell counts were performed in triplicate daily, including immediately upon inoculation, with the MUSE® Cell Analyzer (EMD Millipore, Billerica, MA) and averaged. Specific growth rates were calculated as ln(x_1_-x_0_)/t, where x_1_ = number of cells at the end of exponential growth, x_0_ = number of cells at the beginning of exponential growth, t = number of exponential growth days.

### Photosynthetic Parameter Measurements

Photosynthetic parameter measurements were performed using a Walz WATER-PAM fluorometer (Heinz Walz GbmH, Eichenring, Germany). The fluorometer was calibrated by using ASW as a blank. Cultures at 1-3×10^6^ cells/mL were dark-adapted for at least 45 minutes prior to measurements. Measurements were performed on 3-4 replicate aliquots from the same culture stocks. Fv/Fm, α, rETR_max_, E_k,_ and NPQ were calculated by the WinControl-3 software (Heinz Walz GbmH, Eichenring, Germany).

### Silica Staining and Microscopy

PDMPO (2-(4-pyridyl)-5-((4-(2-dimethylaminoethylaminocarbamoyl)methoxy)phenyl)oxazole), a fluorescent dye that binds to freshly incorporated silica [Shimizu et al. 2001], was used to monitor cell wall formation. 5 mL aliquots of exponentially growing cultures were incubated with .125 µM PDMPO in 40 mL glass culture tubes under the cultivation conditions for 24 hours, allowing for approximately two cell doublings. Cells were imaged using a Zeiss Axio Observer Z1 Inverted Microscope (Carl Zeiss Microimaging Inc., USA). The Zeiss#05 (Ex 395–440 nm, FT 460 nm, Em 470nm LP) filter was used for chlorophyll autofluorescence and Zeiss #21HE (Ex 387/15 nm, FT 409, Em 510/90 nm) was used for PDMPO. Images were acquired using a 40x objective and processed with the AxioVision 4.7.2 software (Carl Zeiss Microimaging Inc., USA).

### Cell/Chloroplast Dimensions and BODIPY Fluorescence

2.5-8.3×10^7^ exponentially growing cells per sample were harvested by centrifugation and stored at −20°C until processing. Pellets were thawed on ice and resuspended in 0.5 mL 2.3% NaCl. 1.3 µL of 1 mg/mL stock of the lipophilic fluorescent dye BODIPY (4,4-difluoro-4-bora-3a,4a-diaza-s-indacene, 493/503, Molecular Probes, ThermoFisher Scientific, USA) per sample was added to stain for neutral lipids (TAG). Following a 30 min incubation on ice in the dark, data for 10,000 cells per sample were collected using an ImageStream X imaging flow cytometer with the INSPIRE^TM^ software package (Amnis Corp., Seattle, WA, USA). 0.6 and 1.0 neutral density filters and 488 nm excitation were used during data aquisition. Post-acquisition spectral compensation and data analysis were performed with the IDEAS^TM^ software (Amnis Corp., Seattle, WA, USA) [Hildebrand et al. 2015]. After debris, unfocused cells, and images containing more than one cell were discarded, 1200-7200 cells were analyzed per sample.

### Protein Content

1.5-7.2×10^7^ cells per replicate were harvested by centrifugation and stored at −20°C until processing. Pellets were thawed on ice and resuspended in 6x volume of extraction buffer (4% SDS, 125 mM Tris-Cl pH 6.8), incubated at 95°C for 5 min, then centrifuged at maximum speed for 3 min. Supernatants were transferred to clean tubes. Protein concentrations were measured using the DC^TM^ Protein Assay (Bio-Rad, Hercules, CA, USA), based on the Lowry method for protein quantification, according to manufacturer’s instructions. Absorbance measurements at 750 nm were performed in triplicate using a SpectraMax M2 microplate reader (Molecular Devices LLC, San Jose, CA, USA). A bovine gamma globulin standard (Bio-Rad, Hercules, CA, USA) set was used to generate a standard curve for calculating protein concentrations in the samples.

### Carbohydrate Content

Total cellular carbohydrate content was determined using a method adapted from Granum and Myklestad [2002]. 1.8-6.3×10^7^ cells per replicate were harvested by centrifugation, washed in 2.3% NaCl, and stored at −20°C until processing. Pellets were thawed on ice, resuspended in 1 mL 0.05 M H_2_SO_4_, incubated in a 60°C water bath for 10 min, then centrifuged at 4000 g for 2 min. Two 400 µL supernatant aliquots per sample were transferred to clean 1.5 mL microfuge tubes for duplicate analysis. 100 µL of 3% freshly prepared aqueous phenol and 1 mL concentrated H_2_SO_4_ were added to each tube, followed by vortexing. After a 30 min incubation at room temperature, samples absorbance at 485 nm was measured in a 1 cm quartz cuvette using a DU^TM^ 730 spectrophotometer (Beckman Coulter, Brea, CA, USA). 155, 38.75, 9.68, and 2.42 mg/L glucose solutions were used to generate a standard curve for calculating carbohydrate concentrations in the samples.

### Statistical Analysis

An online one-way ANOVA calculator (https://www.socscistatistics.com/tests/anova/default2.aspx) was used to assess the statistical significance of the difference or lack thereof between measurements obtained for TG lines and WT cultures. An online Pearson correlation coefficient calculator (https://www.socscistatistics.com/tests/pearson/Default2.aspx) was used to assess two-variable correlations.

## AUTHOR CONTRIBUTIONS

Conception and design (OG, MH), experimental work (OG, DPY), data analysis and interpretation (OG), drafting of the manuscript (OG), critical revision for important intellectual content and final approval (OG, DPY). MH is deceased.

## FUNDING

This work was funded by the US Department of Energy, award DE-EE007689.

---

**Fig. S1.**
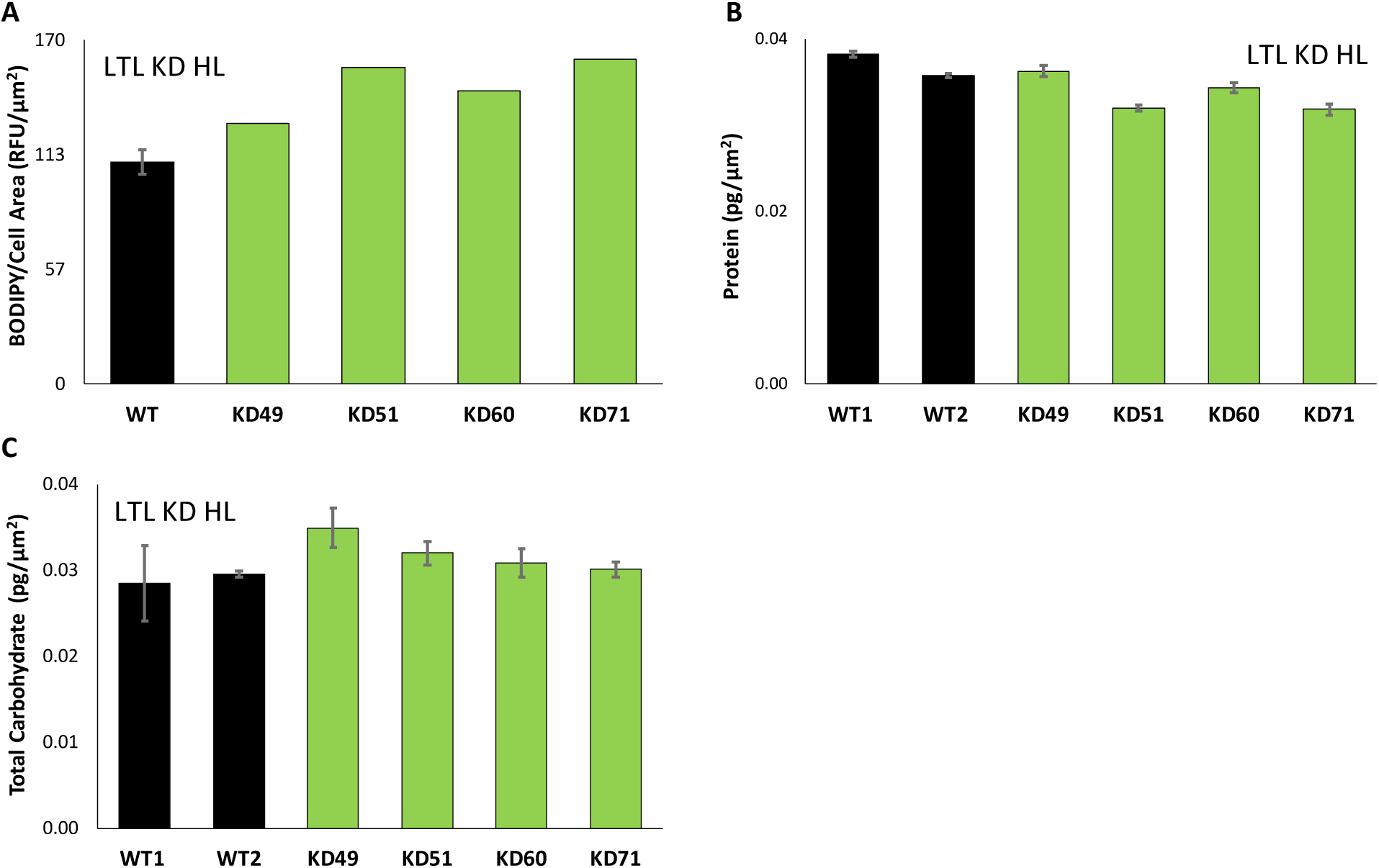
**A.** Average BODIPY fluorescence (relative fluorescence units – RFU) normalized by average cell area. 300 µmol photons m^−2^ sec^−1^ (high light - HL)-adapted LUT1-like knockdown (LTL KD) clones vs. wild-type (WT). WT data is presented as an average of two independent cultures ± standard deviation; **B.** Average cellular protein content normalized by average cell area. HL-adapted LTL KD clones vs. WT. Data are presented as averages of two technical replicates ± standard deviation. **C.** Average total cellular carbohydrate content normalized by average cell area. HL-adapted LTL KD clones vs. WT. Data are presented as averages of two technical replicates ± standard deviation.

**Fig. S2.**
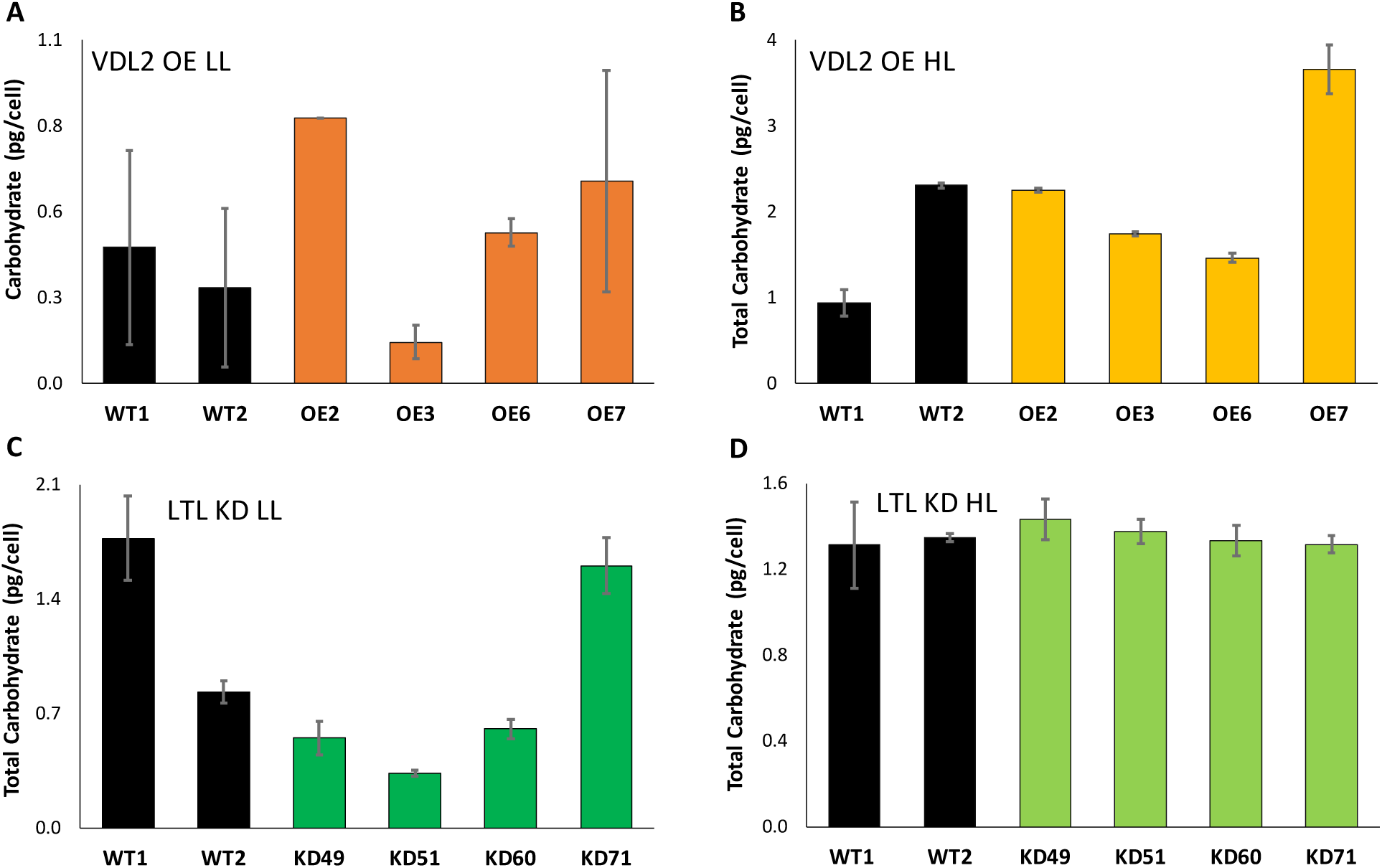
Average total cellular carbohydrate content. Data are presented as averages of two technical replicates ± standard deviation. **A.** 30 µmol photons m^−2^ sec^−1^ (low light - LL)-adapted violaxanthin de-epoxidase-like 2 overexpression (VDL2 OE) clones vs. wild-type (WT); **B.** 300 µmol photons m^−2^ sec^−1^ (high light - HL)-adapted VDL2 OE clones vs. WT; **C.** LL-adapted LUT1-like knockdown (LTL KD) clones vs. WT; **D.** HL-adapted LTL KD clones vs. WT.

**Fig. S3.**
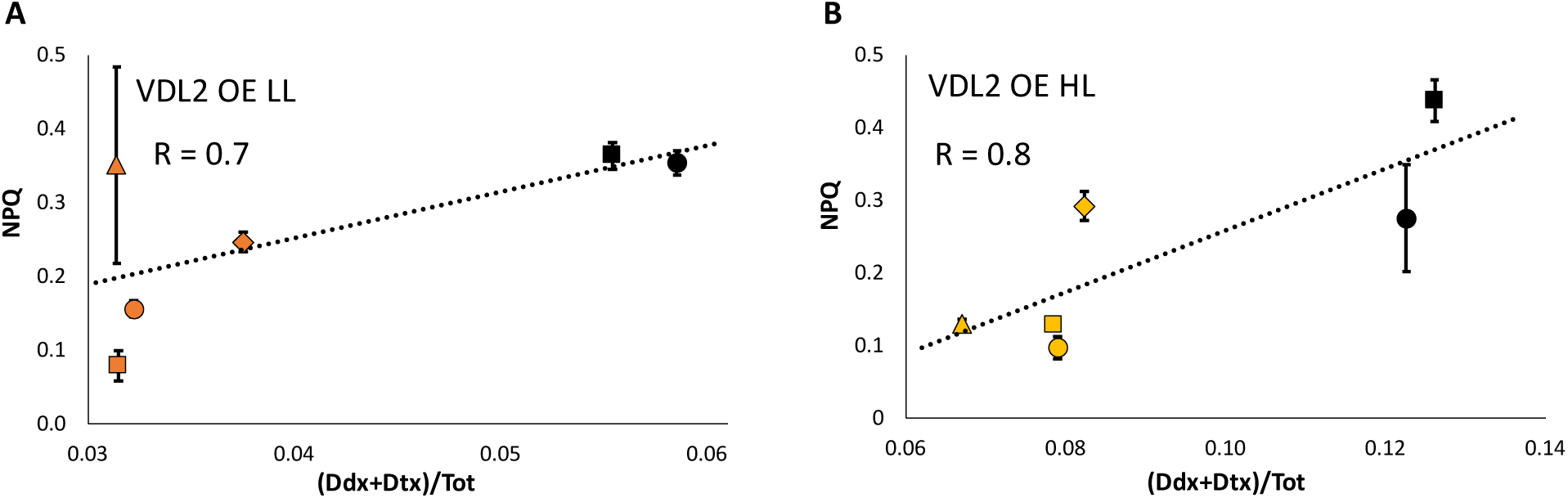
Relationship between the ratio of diadinoxanthin cycle pigments to total cellular photosynthetic pigment content ((Ddx+Dtx)/Tot) [Gaidarenko et al. 2020] and non-photochemical quenching (NPQ) measured at the irradiance closest to cultivation conditions. NPQ values are presented as averages of 3-4 replicates ± standard deviation. Pearson correlation coefficient (R) is indicated on the plots. Wild-type (WT) cultures are represented by black circles (WT1) and squares (WT2). Violaxanthin de-epoxidase-like 2 overexpression (VDL2 OE) clones are represented by orange circles (VDL2 OE2), squares (VDL2 OE3), rhombuses (VDL2 OE6), and triangles (VDL 2 OE7). **A.** 30 µmol photons m^−2^ sec^−1^ (low light - LL)-adapted VDL2 OE clones vs. WT, NPQ measured at 125 µmol photons m^−2^ sec^−1^; **B.** 300 µmol photons m^−2^ sec^−1^ (high light - HL)-adapted VDL2 OE clones vs. WT, NPQ measured at 282 µmol photons m^−2^ sec^−1^.

